# A test of long-term and dose-dependent effects of fluoxetine exposure on the velocity of the invasive crayfish *Procambarus clarkii*

**DOI:** 10.1101/2025.04.11.648412

**Authors:** Daniel Oliveira, Filipe Banha, Pedro Anastácio, Marta C Soares

## Abstract

Fluoxetine, a selective serotonin reuptake inhibitor (SSRI), is one of the most detected antidepressants in wastewater, managing to enter watersheds where it is taken up by freshwater fauna. Here we asked if the serotonergic system affects the dispersal capabilities of *Procambarus clarkii*, a prolific worldwide invasive freshwater crayfish species, thanks in part due to its dispersal rates. For this, we exposed adult crayfish to fluoxetine (serotonin facilitator) and para-chlorophenylalanine (PCPA, serotonin inhibitor) at two concentrations, and measured their velocity after 8 and 16 days, on a two-meter-long metal gutter. Overall, our treatment effects revealed to be non-significant, however the lowest dosage of fluoxetine seem to decrease crayfish mean velocity between the 8th and 16th day of exposure, thus shedding some light on the putative importance of the long-term exposure to environment dosage-dependent fluoxetine in modulating the dispersal capabilities of *P. clarkii*. Though further research is needed, these results can help us better understand the impact of ambient fluoxetine to this invasive species.

## 1. Introduction

Fluoxetine is one of the most prescribed selective serotonin reuptake inhibitors (SSRIs) in western countries. Similarly to other drugs that are excreted or discarded by humans, fluoxetine passes unchanged through wastewater treatments and enters natural waterways (Tierney et al., 2016) making it historically the most detected antidepressant in wastewaters (Fong & Ford, 2014), reaching concentrations as high as 330 ng/L (Mole & Brooks, 2019). Fluoxetine acts acutely by elevating the levels of the neuromodulator serotonin or 5-hydroxytryptamine (5-HT) in synapses. The neuro-endocrine system is evolutionarily conserved both in vertebrates and invertebrates (Tierney et al., 2016) and modulates several functions, such as reproduction, growth, immune function, metabolism, and behaviour (Cheng et al., 2006; Diwan, 2005; Fingerman, 1987, 1997b, 1997a; Fingerman et al., 1981; Huber, Orzeszyna, et al., 1997; Huber, Smith, et al., 1997; Li et al., 2005).

Ambient fluoxetine has been shown to be taken up and stored in animal tissues (Meredith-Williams et al., 2012; Paterson & Metcalfe, 2008). When chronically exposed to, fluoxetine can elicit changes in the release of serotonin, the expression of serotonin receptor genes, and alter the activity of other transmitter systems (Andrews et al., 2015; Kroeze et al., 2012). These changes can cause behavioural changes in many aquatic species, such as fish (e.g. (Correia et al., 2023; Duarte et al., 2019; Eisenreich et al., 2017; Meijide et al., 2018; Pelli & Connaughton, 2015; Vera-Chang et al., 2018; Winder et al., 2012) and crustaceans (e.g.: Bossus, Guler, Short, Morrison, & Ford, 2014; De Lange, Noordoven, Murk, Lürling, & Peeters, 2006; Mesquita et al., 2011).

One important variable that fluoxetine may affect is an organism locomotion (Bossus et al., 2014; De Lange et al., 2006; Mesquita et al., 2011). In fish species, exposure to fluoxetine can decrease swimming activity. For example, Duarte et al. (2019) observed a decrease in *Pomatoschistus microps* active time and an increase in movement delay at concentrations of 1 and 5 mg/L and Meijide et al. (2018) observed that *Gambusia holbrooki* had longer motionless periods at concentrations of 25 μg/L and 50 μg/L. Similar observations were made in crustaceans, with Tierney et al. (2016) observing that juvenile *Orconectes rusticus* exposed for 7 days to concentrations of 2 and 500 μg/L of fluoxetine dissolved in water reduced their locomotion compared to controls. However, fluoxetine can also have the opposite effect, increasing swimming activity. For example, Barry’s (2013) observations reported an increase in swimming velocity in *Aphanius dispar* juveniles exposed to 0.03 and 0.3 μg/L fluoxetine for 7 days. Also, Pan et al. (2018) observed an increase in swimming distance and velocity in *Danio rerio* larvae exposed to 100 μg/L of fluoxetine for 5 days. Additionally, Vera-Chang et al. (2018) reported an increase in the locomotor activity of adult *Danio rerio* exposed to 0.54 μg/L for 6 days. Similarly in crustaceans, Mesquita et al. (2011) exposing *Carcinus maenas* to concentrations of fluoxetine greater than 120 μg/L for 10 days showed that locomotion was significantly increased with animals moving for longer periods of time and walking further.

Similarly to other aquatic species, invasive species locomotion might also be affected by fluoxetine, but no studies currently address this critical issue. This gap is significant since the impact and success of invasive species often depend on their dispersal ability and subsequent ecological effects (Simberloff et al., 2013). One of these species is the red swamp crayfish *Procambarus clarkii*, an omnivore freshwater crayfish of the Cambaridae family, native to northeastern Mexico and the southern USA. This polytrophic keystone species can exert multiple pressures on ecosystems and is a prominent example of a successful invasive species with strong individual variations in dispersal distances (Anastácio et al., 2015; Souty-Grosset et al., 2016) having been introduced in all continents apart from Antarctica and Oceania. It is currently recorded in 16 European countries (Souty-Grosset et al., 2016) and it is considered the most ubiquitous freshwater crayfish species in the world (Chucholl, 2011; Gherardi, 2006; Gutiérrez-Yurrita et al., 2017; Lindqvist & Huner, 2017). It’s success as an invasive species stems from characteristics such as early maturity, fast growth rate, high offspring number, rapid and adaptable life cycle, flexible feeding strategy, and a wide ability to disperse and to tolerate extreme environmental changes (Alcorlo et al., 2004; Banha & Anastácio, 2014; de Abreu et al., 2020; Gherardi, 2007; Gutiérrez-Yurrita et al., 2017; Hobbs et al., 1989; Loureiro et al., 2015; Souty-Grosset et al., 2006, 2016) such as in temporary streams (for ex.: in southern Portugal; Gherardi et al., 2002) and polluted habitats (Gherardi & Barbaresi, 2000).

Due to its invasive nature, understanding of how *P. clarkii* dispersion may be modulated by the serotonergic system can prove important to predict its potential spread and help formulate new plans and strategies to limit its dispersion. As such, the aim of this study is to test the effects of exogeneous fluoxetine exposure on crayfish locomotion. To achieve this, we measured the velocities of *P. clarkii* individuals exposed for 16 days to two concentrations of fluoxetine as well as para-chlorophenylalanine (PCPA), a tryptophan hydroxylase inhibitor that acts as a potent depletor of tissue serotonin in animals (Maximino et al., 2013) to also test the effects of lowering serotonin levels. Considering previous studies on crayfish (ex.: Tierney et al. (2016) as well as other on teleost fish species (ex.: Duarte et al., 2019; Meijide et al., 2018), we expected that exposure to fluoxetine would lead to crayfish velocity decrease, whereas we expected it to increase with PCPA exposure, due to its antagonistic nature to fluoxetine.

## 2. Materials and Methods

### 2.1 Crayfish capture and housing

*Procambarus clarkii* individuals were captured in the marateca stream next to rice fields near the town of Cabrela, Portugal (38.58561 N, 8.528580 W). These were captured with crayfish traps baited with sardines, on two separate occasions: between the 11th to 12th of May 2023 and between the 13th to 14th of July 2023. These catches yielded only adults, with no individual below 25 mm of cephalothorax length. This was most likely due to trap catches being biased towards capturing larger individuals (Dorn et al., 2005).

After capture, crayfish were maintained at 20°C in tanks filled with a depth of 5 to 7 cm of tap water aerated using an aquarium pump. Several PVC tube sections were scattered on the bottom of the tanks to serve as shelter and individuals were fed with carrots. Every four to seven days crayfish were removed temporarily to clean the containers, replace the water and replenish food items.

Crayfish wet weight and cephalothorax length were measured to form groups with crayfish of similar size and weight. Crayfish condition was calculated using Fulton’s condition index with the formula K = W/L^3^, where W is wet weight (g) and L is Cephalothorax length (mm) (Ricker, 1975). The selected crayfish were then stored individually inside small containers with 300 ml of internal volume for a minimum of 5 days before the experiment. The top-covers of the containers were perforated, and an appropriate volume of open-air space was given to allow proper oxygenation of the water. The containers were cleaned and the water and uneaten food replaced with fresh water and food every two days.

### 2.2 Experimental design and physiological treatments

The general set-up consisted of a two-meter-long metal gutter filled with a depth of 7 cm of tap water that was replaced a day before every experiment. A substrate consisting of medium coarse sand with approximately 1cm of dept was used to cover the metal gutter’s bottom. A removable translucent plastic plate separated the initial 10 cm of the track from the rest of it.

We used 144 adult crayfish in this experiment, with a mean cephalothorax length of 45.01 mm (sd = 3.67 mm). There were 85 males and 59 females, seven of which were carrying eggs. These crayfish were grouped into 5 treatments balanced for size. Each group was immersed in 200 ml of solutions of either 1μg/L of fluoxetine, 100μg/L of fluoxetine, 1μg/L of PCPA or 100μg/L of PCPA. The fifth group acted as a control and was immersed in 200 ml of regular tap water. The mean cephalothorax length for the fluoxetine 1μg/L was 45.04 mm (sd = 3.758), for the fluoxetine 100μg/L it was 45.00 mm (sd = 3.332), for the PCPA 1μg/L it was 44.98 mm (sd = 4.15), for the PCPA 100μg/L it was 44.97 (sd = 3.766) and for the control group it was 45.04 mm (sd = 3.398). The lowest concentration used, which was higher than the maximum freshwater levels of fluoxetine detected (Mole & Brooks, 2019), was chosen in the expectation that an affect might be observed. The 100 μg/L was chosen as to give a noticeably distinct order of magnitude. PCPA at the same concentrations was used to test the effects of serotonin disruption. The solutions and food items were replaced with new solutions and food every three days. Crayfish were exposed to the solutions for 16 days, with experiments conducted on the 8th and 16th days, giving a total of two days of experiments.

### 2.3 Behavioural analysis

Crayfish went through 3 days of fasting and were acclimatised to a temperature of 22°C for 24 hours before the day of the experiment. The temperature used corresponds to water temperatures registered in spring and summer, respectively, in rivers from the south of Portugal (Anastácio et al., 1999; Reis & Araujo, 2016). The pH, percentage and ppm of dissolved oxygen and temperature were recorded before and after the experiments to ensure consistency through the two days of experimentation. For each treatment, an individual was placed on the first 10 cm of the track and was kept for 60 seconds from roaming by the closed divider, to acclimatize to the new environment. After 60 seconds, the divider was removed to allow the individual to roam freely for 120 seconds or until it reached the opposite end of the track, after which its position or the time it took to reach the end of the gutter was noted. The dispersal capability of individuals was assessed by calculating their mean velocity.

### 2.4 Statistical analysis

All statistical analysis were done using RStudio (2023.06.1 Build 524, Posit Software, PBC) with the libraries tidyverse, car, lme4, rstatix and fitdistrplus.

Velocity values underwent a log(x+1) transformation in order to normalize the data and fulfil the homoscedasticity assumption required for parametric statistical analysis (function leveneTest() from the car package). Sphericity was also tested.

A repeated measures ANOVA (function anova_test() from the rstatix package) was used, followed by a pairwise t-test (function pairwise_t_test() from the rstatix package) to determine if there were differences in the treatments between the two days of experimentation.

## 3. Results

Overall, crayfish velocity was not significantly different across the various treatments (Repeated measures ANOVA: F= 2.183; p=0.071). Velocity was also not different between sexes (Two Sample t-test: t = 1.218; p-value = 0.224).

Interestingly, pairwise t-tests for crayfish velocities revealed that those treated with fluoxetine 1μg/L decreased in speed between the 8th and 16th day, with an average velocity of 1.296 cm/s (sd = 1.036) on the 8th day and 1.0542 cm/s (sd = 1.052) (see Figure 1 and Table 1). No other significant results were found (see Table 1).

**Table 1.** Pairwise t-test results for velocity on the 8th and 16th day of exposure. n1 and n2 correspond to the sample counts, statistic values correspond to the values used to compute the p-value and df correspond to the degrees of freedom. Bonferroni-adjusted p values are presented.

| Pair-Wise t-test | Velocity |  |  |  |  |  |  |  |  |  |
| --- | --- | --- | --- | --- | --- | --- | --- | --- | --- | --- |
|  | Treatment | .y. | group1 | group2 | n1 | n2 | statistic | df | p | Sig. |
|  | Control | Vel. | 8 | 16 | 29 | 29 | -1.495 | 28 | 0.146 | ns |
|  | Fluoxetine1μg | Vel. | 8 | 16 | 28 | 28 | 2.617 | 27 | 0.014 | * |
|  | Fluoxetine 100μg | Vel. | 8 | 16 | 29 | 29 | -1.137 | 28 | 0.265 | ns |
|  | PCPA 1μg | Vel. | 8 | 16 | 29 | 29 | -0.428 | 28 | 0.672 | ns |
|  | PCPA 100μg | Vel. | 8 | 16 | 29 | 29 | -0.794 | 28 | 0.434 | ns |

**Figure 1.**
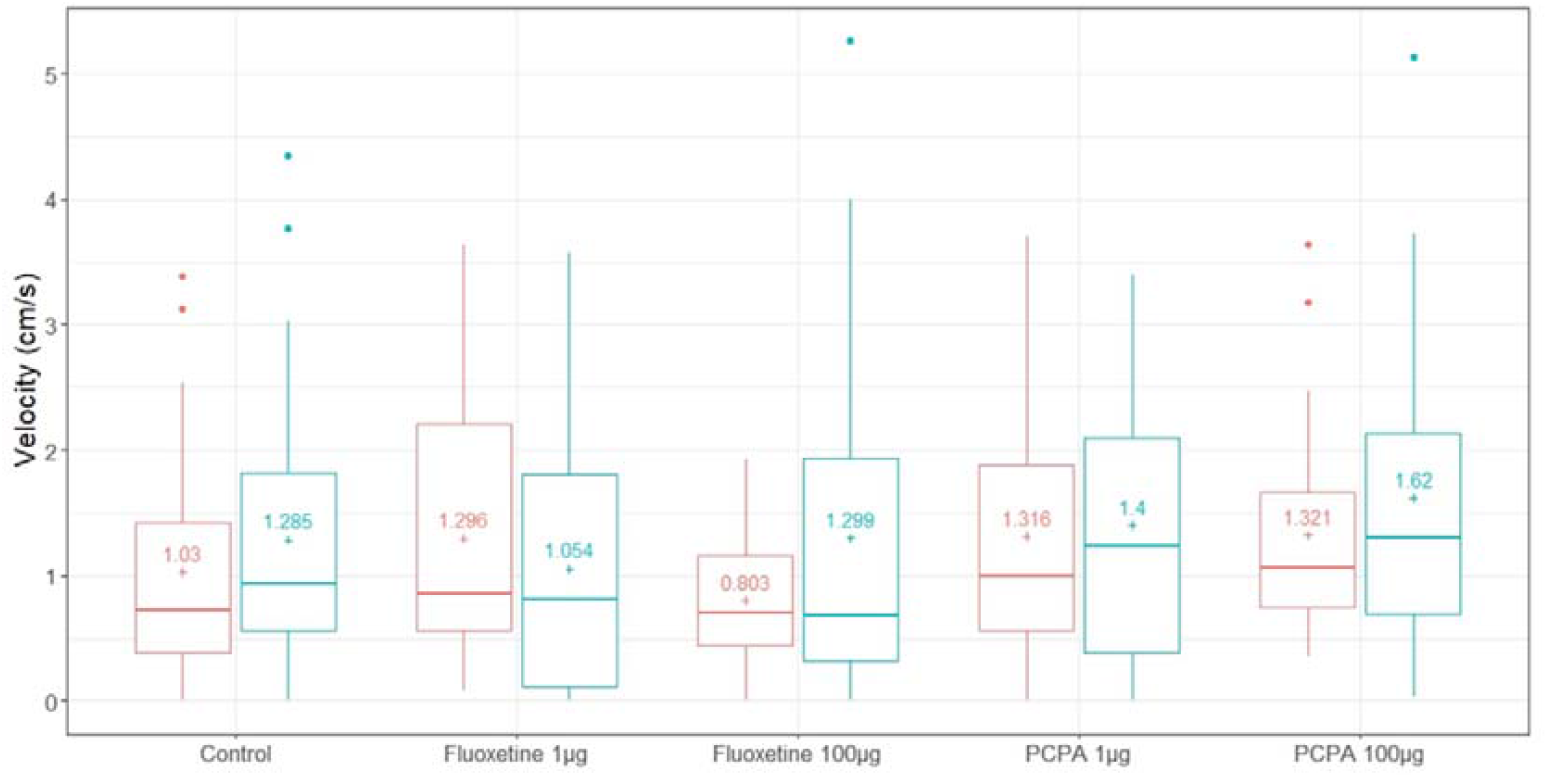
Boxplot displaying variation of velocity between treatments on the 8^th^ and 16^th^ days of experiment. Left/red coloured plots represent the 8^th^ day and right/blue coloured plots represent the 16^th^ day. Plus sign indicates the mean value.

### 4. Discussion

The role of fluoxetine, and by extension that of serotonin, on locomotion can be of great importance to ecology, especially concerning how exogenous chemical compounds may modulate the dispersion of invasive species. The prediction of which species are candidate invaders has been a long-standing dilemma and of increasing interest to ecologists (Forsyth et al., 2004; Kolar & Lodge, 2001) and their ability to disperse may be a key factor in determining the successes of invasion as well as the impact and rate of spread (Ehrlich, 1989; Lodge, 1993). Overall, we found that the velocity values did not differ significantly between the treatments and control group, which opposes previous studies on fish that obtained noticeable differences with concentrations of fluoxetine, comparable to those used in this study. For example, Meijide et al. (2018) observed a decrease in swimming behaviour of *Gambusia holbrooki* at concentrations of 25 μg/L and 50 μg/L and Pan et al. (2018) reported increase in distance travelled and faster swimming speeds in *Danio rerio* larvae exposed to 100 μg/L of fluoxetine. However, *P. clarkii* is known to possess remarkable tolerance to polluted environments, being capable of accumulating heavy metals and toxins (Suárez-Serrano et al., 2010) and to have higher resistance to drugs such as maduramicin (Gao et al., 2021). Moreover, antagonists, such as PCPA, generally need to be administered in larger doses that exceed what is normally required to overcome the agonist effects of the biomolecule they are trying to counteract (Muir, 2015). Because *P. clarkii* may seem to show a greater resilience to fluoxetine and PCPA, putatively requiring higher or longer exposures to cause immediate noticeable alterations to their locomotion, dosages used in our study may have been too low to elicit more conspicuous responses. Indeed, some studies done on other crustacean species reported fluoxetine influencing locomotion at concentrations higher than those used in this study (ex.: Mesquita et al., 2011; Tierney et al., 2016). Another important variable is the fact that our used individuals were all adults, which may deemed them even more resistant to serotonergic drugs. Tierney et al. (2016) observed that adult *Orconectes rusticus* walking behaviour did not differ significantly from controls when these were exposed to fluoxetine, which led the author to suggest that adult crayfish are more resistant to the behavioural effects of fluoxetine than juveniles are. This too could be the case for *P. clarkii*.

The effects of lower dosage fluoxetine (1μg/L) treatment were particularly interesting between the 8th and 16th days of experimentation, with the average velocities decreasing during the study. This points towards different impacts of SSRI in accordance with dosage (ambient concentrations) and time of exposure. For instance, studies have found that either decreases or increases in some behavioural variables such as aggression in crustaceans exposed to serotonin or SSRIs are dependent on the dose and timescale of exposure (Reisinger et al., 2021). This may occur due to stress mediation, which in crayfish happens in connection with the serotonergic system (Fossat et al., 2014, 2015). For instance, Fossat and colleagues (2014) showed that stressed crayfish (*Procambarus clarkii*) expressed anxiety-like behaviour that was modulated by 5HT. In our case, the dose dependent rise of fluoxetine may have increased individuals stress levels, which may impacted their mobility over the 15 days of exposure. In a greater dosage of fluoxetine, the putative more rapid increase of stress could have led to a shutting down response (aka negative feedback), which may have physiological and immune consequences, but these were not distinguishable by their mobility.

Fluoxetine mobility effects seem to depend on the species tested. Most of the studies found reductions on distances travelled, the number of times they started moving as well as delays in starting to move (ex.: Duarte et al., 2019; Eisenreich et al., 2017; Meijide et al., 2018), while a few report an increase in locomotor activity (ex.: Barry, 2013; Pan et al., 2018; Vera-Chang et al., 2018). Though the exposure period is somewhat comparable to previous studies (Mesquita et al., 2011; Tierney et al., 2016), and considering the potential resilience of *P. clarkii*, this exposure period may have not been long enough for fluoxetine to elicit a more significant response. A longer exposure period, possibly measured in months rather than weeks, could elicit even more noticeable effects.

Our findings suggest that, overall, these drugs do not alter *P. clarkii* locomotion in the tested exposure periods and doses, but it does shed some light on the putative importance of the long-term exposure to environment dosage-dependent fluoxetine in modulating the dispersal capabilities of *P. clarkii*. Because these mobility-related changes were not occurring overall and were just seen in a particular dose (the smaller one) it suggests that exposure time is important, but also that animals’ brains respond differently to variable exposures. Faced with current global changes and habitat alterations it is important to continuously expand our knowledge and understanding of how the presence of drugs such as fluoxetine affect the biology of aquatic organisms. Doing so can prove instrumental in formulating new plans and strategies to mitigate the impacts of invasive species such as *P. clarkii*.

## 6. Acknowledgements

This study was financially supported by the LIFE INVASAQUA project (Aquatic Invasive Alien Species of Freshwater and Estuarine Systems: Awareness and Prevention in the Iberian Peninsula) (LIFE17 GIE/ES/000515) funded by the EU LIFE program and supported by the strategic plan of MARE – Marine and Environmental Sciences Centre (UID/MAR/04292/2019). M.C.S. and F.B. were supported by the Foundation for Science and Technology through an individual contract (2021.01458.CEECIND/CP1668/CT0003 and CEEC/01896/2021, respectively).

## 7. Author contributions

M.C.S. and F.B. designed the study. D.O. and F.B. collected the animals. D.O. collected and analysed the data. D.O. wrote first version of the manuscript. M.C.S, F.B. P.A. and D.O. reviewed and edited all versions of the manuscript.

## Annexes

**Figure A1.**
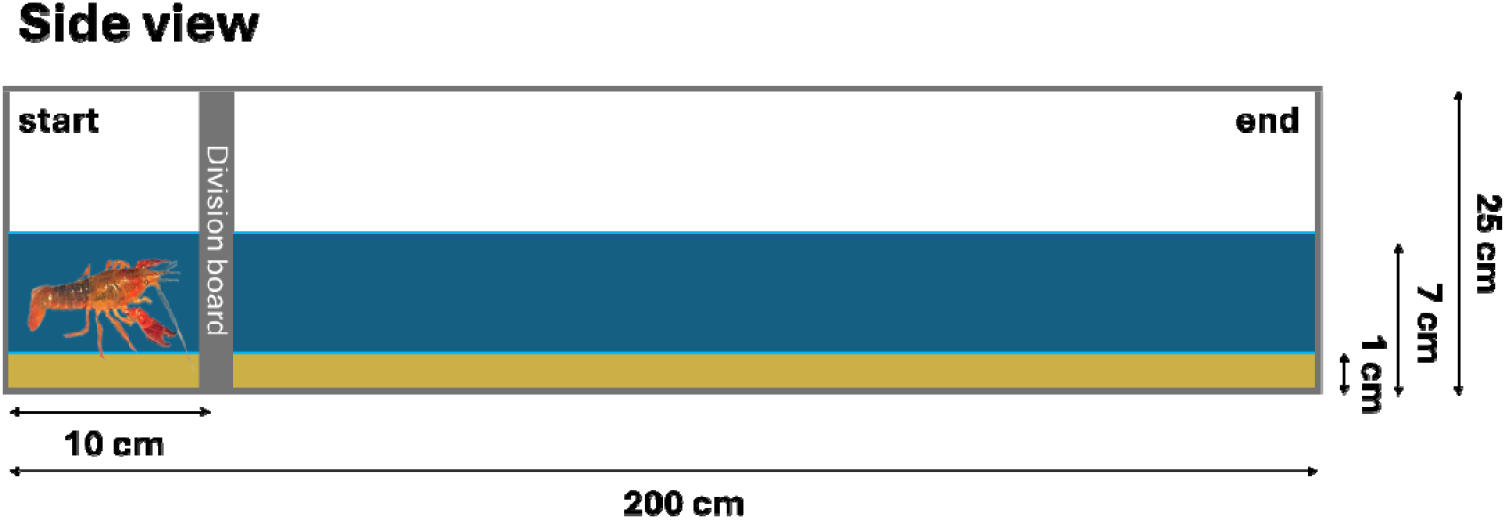
Schematic side view of the general layout used for the experiments

